## Supplementary material for "The evolution of skilled hindlimb movements in birds: A citizen science approach": Suplementary Materials

**Supplementary Materials**

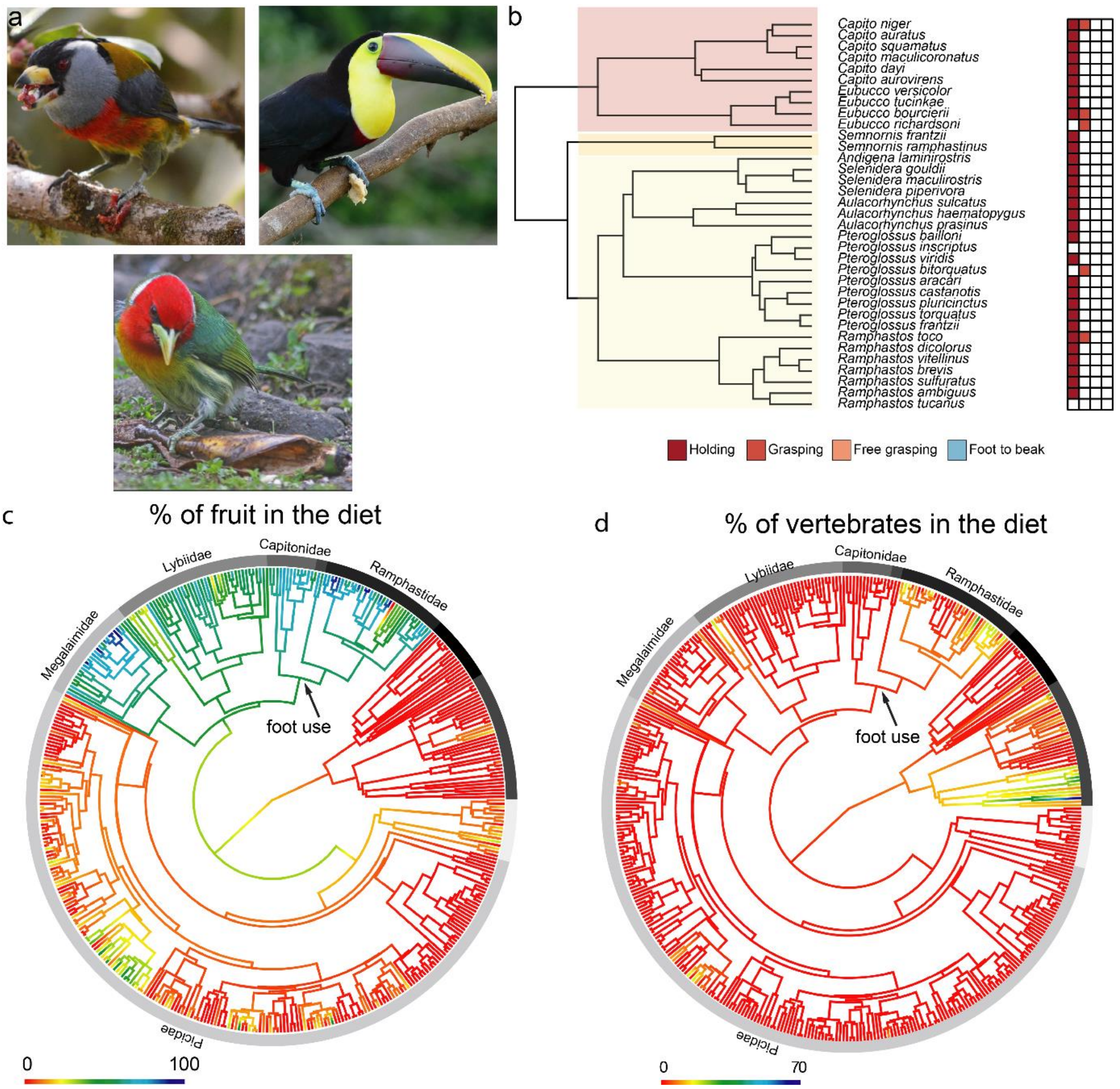


**Supplementary figure 1. Foot use evolution in new world barbets and toucans. a,** examples of foot use in the three families of Piciformes where foot use is present, Semnornithidae (Toucan-Barbets. Top left: Toucan Barbet, *Semnornis ramphastinus*), Rhamphastidae (Toucans and toucanets, top righ: Yellow-throated Toucan, *Ramphastos ambiguous*), and Capitonidae (New world barbets, bottom panel: Red-headed Barbet, *Eubucco bourcierii*). Photographer credits are listed in Table S7. **b,** a character matrix species level phylogenies for the same three families. Grasping is rare (only recorded in five species) and most species hold objects against a perch. **c** and **d** show ancestral state reconstruction for all species of Piciformes with the percentage of the diet that is constituted by fruits (c) and vertebrates (d) mapped. The node where foot use is likely to have evolved is indicated.

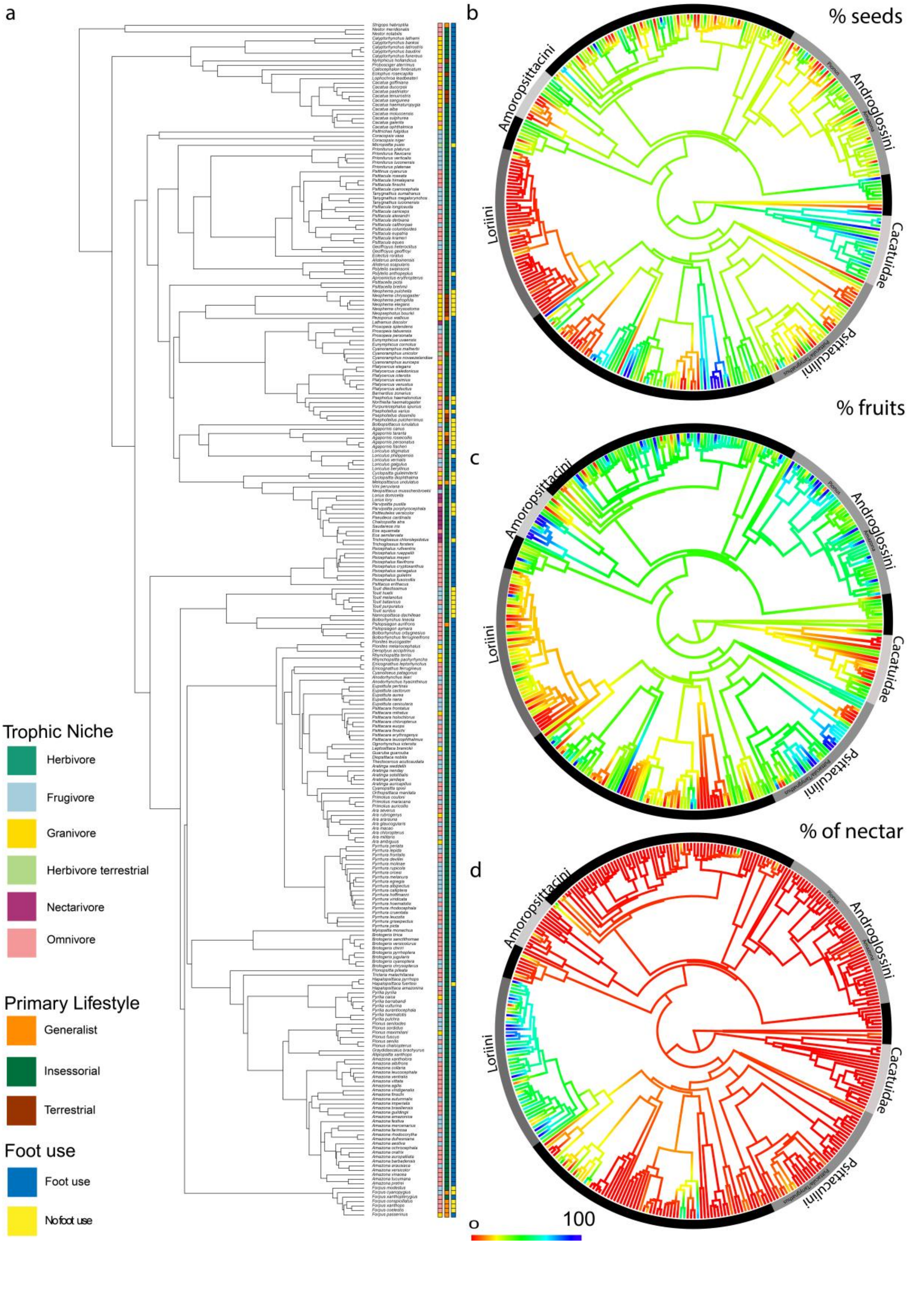
a
b
% seeds
c
% fruits
d
% of nectar
Amoropsittacini
Androglossini
Loriini
Cacatuidae
Psittaculini
0
100
Trophic Niche
Herbivore
Frugivore
Granivore
Herbivore terrestrial
Nectarivore
Omnivore
Primary Lifestyle
Generalist
Insessorial
Terrestrial
Foot use
Foot use
No foot use

**Supplementary figure 2. Interplay between ecological and trophic characteristics and foot use among parrots.** A, character matrix showing habitat, trophic niche, primary lifestyle and foot use in 282 species of parrots. **b**, **c**, and **d**, show ancestral state reconstruction for all species of parrots of the percentage of the diet that is constituted by seeds (b), fruits (c) and nectar. (d).

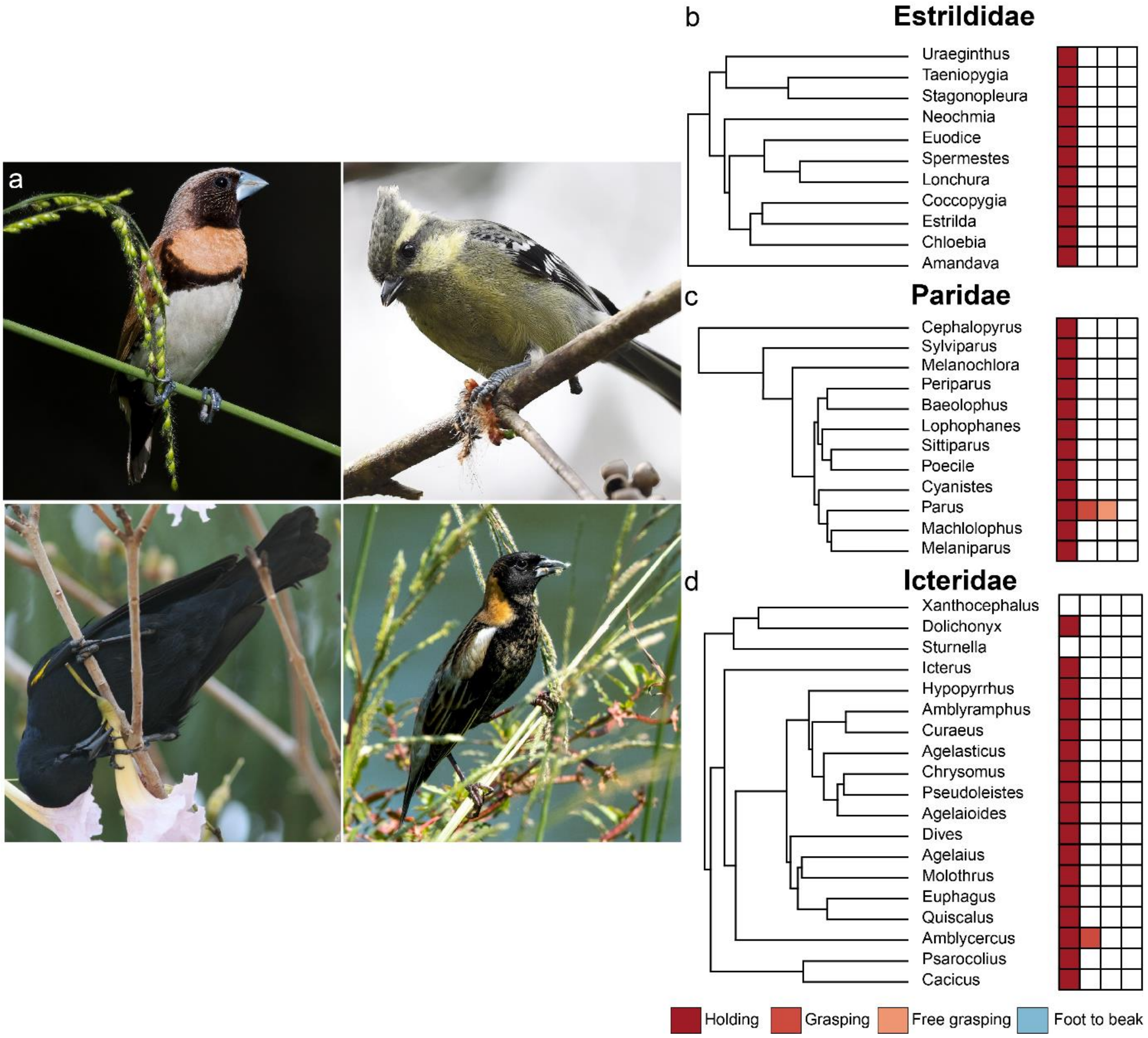


**Supplementary figure 3: foot use three families of Passeriformes**. **a**, shows examples of members of the families estrildid (top left: Chestnut-breasted Munia, *Lonchura castaneothorax*), Paridae (top right: Indian Yellow Tit, *Machlolophus aplonotus*) and Icteridae (bottom two panels, left: Yellow-shouldered Blackbird, *Agelaius xanthomus*; right: Bobolink, *Dolichonyx oryzivorus*) using their feet to manipulate objects. Photographer credits are listed in Table S7. **b, c** and **d** show a character matrix at the genus level for the same families. colored squares reflect the presence of each of the four behavioral elements.

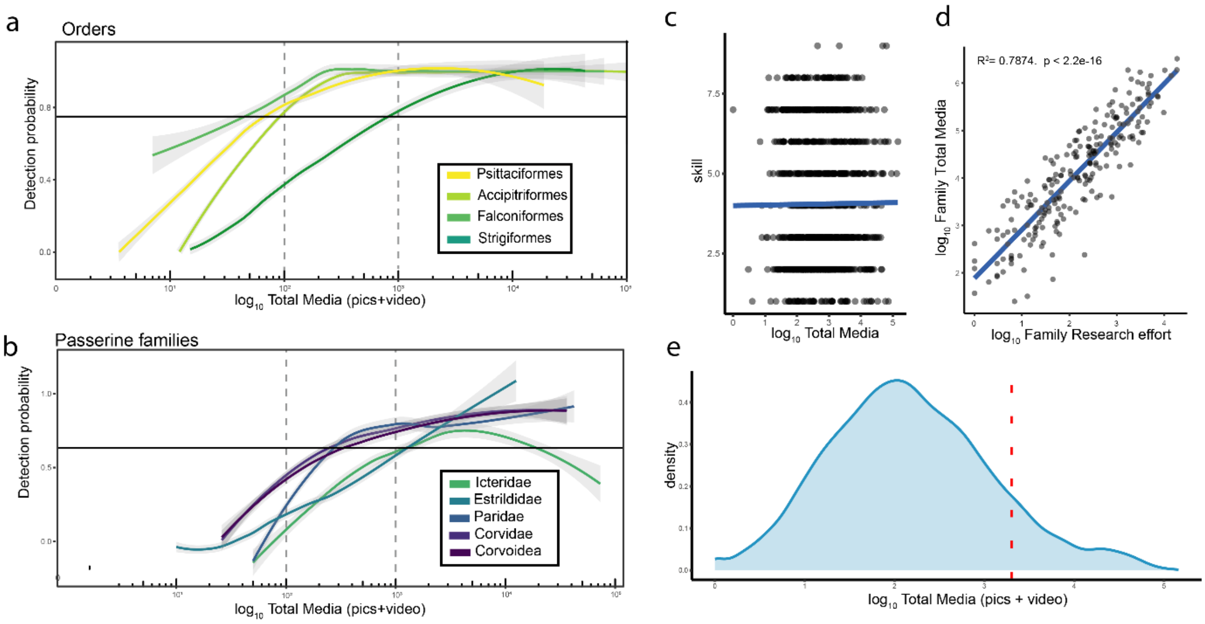


**Supplementary figure 4. a, and b,** show the correlation between total media and the probability of detecting foot use behavior for different orders (a) and songbird families (b). The curves were used to determine the threshold at which foot use behavior could be detected for each clade. Dotted lines show where the lines interest with 100 and 1000 pictures. Solid lines show the 75% detection probability. **c**, Foot use skill score plotted against log-transformed total media available for 1020 species of birds. Across all birds we found no correlation between skill scores and media available (PGLS, $F_{1,1018,}$= 2.877, p= 0.26). Blue line shows PGLS regression line. **d,** a scatterplot of total media available in the Macaulay Library at the family level plotted against research effort, defined as the total number of papers published on each family ((Ducatez et al., 2020)). The solid line indicates a significant correlation between the two (PGLS: $F_{1,202,}$= 103.4, p= >0.001). **e**, A density plot showing the distribution of the total number of media for all species of birds in the Macaulay Library. The red line show where 90 % of the species are found.

**Table S1: Model parameters for hidden rates analysis for family level foot use behavior.**

| | *-lnL* | *AIC* | *AICc* | *# Rate cat* |
|---|---|---|---|---|
| *ER* | -141.47 | 284.94 | 284.95 | 1 |
| *SYM* | -141.47 | 284.94 | 284.95 | 1 |
| *ARD* | -133.27 | 270.55 | 270.60 | 1 |
| *PREC* | -127.92 | 259.84 | 259.89 | 2 |

-lnL=maximum log-likelihood; AIC=Akaike information criterion; AICc=Akaike information criterion corrected for sample size. ER = equal rates, SYM= symmetrical model, all-rates different matrix model, PREC = precursor model. # Rate cat = number of rate categories

**Table S2. Ancestral Diets. Ancestral diet reconstruction.** Numbers indicate the ancestral state probability for the most recent common ancestor for each clade (see methods for details).

| *Clade* | *Aquatic pred.* | *Frugivore* | *Granivore* | *Herbivore* | *Invertivore* | *Nectarivore* | *Omnivore* | *Vertivore* |
|---|---|---|---|---|---|---|---|---|
| *Accipitriformes* | 0.000 | 0.000 | 0.000 | 0.001 | 0.000 | 0.000 | 0.000 | 0.999 |
| *Strigiformes* | 0.000 | 0.000 | 0.000 | 0.001 | 0.000 | 0.000 | 0.000 | 0.999 |
| *Falconiformes* | 0.000 | 0.000 | 0.000 | 0.001 | 0.000 | 0.000 | 0.000 | 0.999 |
| *Psittaciformes* | 0.003 | 0.015 | 0.694 | 0.063 | 0.001 | 0.001 | 0.210 | 0.012 |
| *Passeriformes* | 0.000 | 0.000 | 0.000 | 0.000 | 1.000 | 0.000 | 0.000 | 0.000 |
| *Passerida* | 0.000 | 0.000 | 0.003 | 0.000 | 0.980 | 0.000 | 0.016 | 0.000 |
| *Corvides* | 0.000 | 0.000 | 0.000 | 0.000 | 1.000 | 0.000 | 0.000 | 0.000 |
| *Piciformes* | 0.000 | 0.004 | 0.000 | 0.000 | 0.995 | 0.000 | 0.000 | 0.000 |
| *Capitonidae, Semnornithidae,* and *Ramphastidae* | 0.000 | 0.998 | 0.000 | 0.000 | 0.000 | 0.000 | 0.002 | 0.000 |
| *Sylvioidea* | 0.00 | 0.00 | 0.00 | 0.00 | 0.87 | 0.00 | 0.13 | 0.00 |

**TableS3. Detection threshold.**

| | ***calc. 75 % detection threshold*** | ***Used threshold*** |
|---|---|---|
| *Accipitriformes* | 108 | 110 |
| *Strigiformes* | 1009 | 1050 |
| *Falconiformes* | 77 | 80 |
| *Psittaciformes* | 66 | 70 |
| *Piciformes* | 573 | 700 |
| *Corvidae* | 338 | 350 |
| *Paridae* | 326 | 330 |
| *Estrildidae* | 1631 | 1700 |
| *Icteridae* | 1822 | 1900 |
| *Corvoidea* | 488 | 500 |

**Table S4: Model parameters for hidden rates analysis for foot use behavior in parrots**

| *model* | *-lnL* | *AIC* | *AICc* | *Rate cat* |
|---|---|---|---|---|
| *ER* | -203 | 408.04 | 408.01 | 1 |
| *SYM* | -194.977 | 401.95 | 402.28 | 1 |
| *ARD* | -174.122 | 372.24 | 373.50 | 1 |

-lnL=maximum log-likelihood; AIC=Akaike information criterion; AICc=Akaike information criterion corrected for sample size. ER = equal rates, SYM= symmetrical model, all-rates different matrix model. # Rate cat = number of rate categories

**Table S5. Pictures information**

| *ML ID* | *Species name* | *English name* | *photographer* | *figure* |
|---|---|---|---|---|
| *ML57273391* | *Melanochlora sultanea* | Sultan Tit | Craig Brelsford | fig. 1 |
| *ML219550691* | *Anodorhynchus hyacinthinus* | Hyacinth Macaw | Nick Athanas | fig. 1 |
| *ML204333801* | *Elanoides forficatus* | Swallow-tailed Kite | Hal and Kirsten Snyder | fig. 1 and 3 |
| *ML126826011* | *Athene cunicularia* | Burrowing Owl | Brad Imhoff | fig. 3 |
| *ML213918601* | *Microhierax caerulescens* | Collared Falconet | Dominic Standing | fig. 3 |
| *ML36360411* | *Accipiter rufitorques* | Fiji Goshawk | Mat Gilfedder | fig. 3 |
| *ML205654991* | *Gypohierax angolensis* | Palm-nut Vulture | Frans Vandewalle | fig. 3 |
| *ML204478221* | *Aegypius monachus* | Cinereous Vulture | Juan Lacruz Martin | fig. 3 |
| *ML205223501* | *Nestor meridionalis* | New Zealand Kaka | Dave Rintoul | fig. 4 |
| *ML66912201* | *Psittacara erythrogenys* | Red-masked Parakeet | Walter Oshiro | fig. 4 |
| *ML131156891* | *Poicephalus cryptoxanthus* | Brown-headed Parrot | Volker Hesse | fig. 4 |
| *ML176135231* | *Amazona aestiva* | Turquoise-fronted Parrot | Estevão Freitas Santos | fig. 4 |
| *ML233718501* | *Cacatua sanguinea* | Little Corella | Michael Daley | fig. 4 |
| *ML293520261* | *Psittacula krameri* | Rose-ringed Parakeet | Dobrin Botev | fig. 4 |
| *ML58505951* | *Cyanocorax yucatanicus* | Yucatan Jay | Chris Wood | fig. 5 |
| *ML449915461* | *Vireo flavifrons* | Yellow-throated Vireo | John Garrett | fig. 5 |
| *ML141977161* | *Mohoua albicilla* | Whitehead | Kuang-Ping Yu | fig. 5 |
| *ML409098921* | *Cracticus nigrogularis* | Pied Butcherbird | Peter Lowe | fig. 5 |
| *ML327037341* | *Dicrurus bracteatus* | Spangled Drongo | Ged Tranter | fig. 5 |
| *ML231781581* | *Lanius minor* | Lesser Gray Shrike | Haldun Savaş | fig. 5 |
| *ML229677561* | *Dolichonyx oryzivorus* | Bobolink | Court Harding | fig. S3 |
| *ML144678231* | *Machlolophus aplonotus* | Indian Yellow Tit | Sriram Reddy | fig. S3 |
| *ML321218961* | *Lonchura castaneothorax* | Chestnut-breasted Munia | Terence Alexander | fig. S3 |
| *ML204717411* | *Agelaius xanthomus* | Yellow-shouldered Blackbird | Mikko Pyhälä | fig. S3 |
| *ML83544481* | *Eubucco bourcierii* | Red-headed Barbet | Larry Therrien | fig. S1 |
| *ML90815121* | *Semnornis ramphastinus* | Toucan Barbet | Eli Gross | fig. S1 |
| *ML79312761* | *Ramphastos ambiguus* | Yellow-throated Toucan | Krista Kaptein | fig. S1 |

**Table S6.** List of species and source of foot use in the literature.

| Orders | Family | English | Scientific name | Source |
|---|---|---|---|---|
| Accipitriformes | Accipitridae | Tawny eagle | Aquila rapax | (Van Someren, 1956) |
| Accipitriformes | Accipitridae | Verreaux's eagle | Aquila verreauxii | (Mundy et al., 1986) |
| Accipitriformes | Accipitridae | Double-toothed kite | Harpagus bidentatus | (Baker et al., 1999) |
| Accipitriformes | Accipitridae | Plumbeous kite | Ictinia plumbea | (Seavy et al., 1998) |
| Accipitriformes | Accipitridae | Bat hawk | Macheiramphus alcinus | (Thomsett, 1981) |
| Accipitriformes | Accipitridae | The yellow-billed kite | Milvus aegyptius | (Finn, 1908; Van Someren, 1956) |
| Accipitriformes | Cathartidae | Turkey vulture | Cathartes aura | (Kempton, 1927) |
| Accipitriformes | Pandionidae | Ospreys | Pandion haliaetus | (Sustaita et al., 2019) |
| Accipitriformes | Sagittariidae | Secretary bird | Sagittarius serpentarius | (Van Someren, 1956) |
| Cariamiformes | Cariamidae | Red-legged seriema | Cariama cristata | (Finn, 1911; Oswald et al., 2023) |
| Charadriiformes | Chionidae | Snowy sheathbill | Chionis albus | (Finn, 1928) |
| Charadriiformes | Haematopodidae | Pied oystercatcher | Haematopus longirostris | (Jones, 1867) |
| Charadriiformes | Stercorariidae | Parasitic jaeger | Stercorarius parasiticus | (Pruett-Jones, 1980) |
| Ciconiiformes | Ciconiidae | Asian openbill stork | Anastomus oscitans | (Bingham, 1876) |
| Coliiformes | Coliidae | White-backed mousebird | Colius colius | (Rowan, 1967) |
| Coliiformes | Coliidae | Red-faced mousebird | Colius indicus | (Rowan, 1967) |
| Coliiformes | Coliidae | Speckled mousebird Birds | Colius striatus | (Rowan, 1967) |
| Columbiformes | Columbidae | Tooth-billed pigeon | Didunculus strigirostris | (Gifford, 1925; Collar, 2015) |
| Columbiformes | Columbidae | Marquesan ground dove | Pampusana rubescens | (Smith, 1971) |
| Cuculiformes | Cuculidae | Greater coucal | Centropus sinensis | (Finn, 1922) |
| Cuculiformes | Cuculidae | Guira cuckoo | Guira guira | (Finn, 1919) |
| Falconiformes | Falconidae | Crested caracara | Caracara plancus | (Kilham, 1979) |
| Galliformes | Megapodiidae | Australian brushturkey | Alectura lathami | (Finn, 1919) |
| Gruiformes | Aramidae | Limpkin | Aramus guarauna | (Billerman et al., 2022) |
| Gruiformes | Rallidae | Weka | Gallirallus australis | (Billerman et al., 2022) |
| Gruiformes | Rallidae | Allen's gallinule | Porphyrio alleni | (Billerman et al., 2022) |
| Gruiformes | Rallidae | Purple gallinule | Porphyrio martinica | (Billerman et al., 2022) |
| Gruiformes | Rallidae | Western swamphen | Porphyrio porphyrio | (Billerman et al., 2022) |
| Passeriformes | Aegithalidae | Long-tailed Tit | Aegithalos caudatus | (Harrap, 2020) |
| Passeriformes | Aegithalidae | Long-tailed Tit | Aegithalos caudatus | (Simmons, 1963) |
| Passeriformes | Aegithinidae | Common iora | Aegithina tiphia | (Finn, 1908) |
| Passeriformes | Artamidae | Lack-faced woodswallow | Artamus cinereus | (Immelmann, 1966) |
| Passeriformes | Artamidae | White-breasted Woodswallow | Artamus leucorynchus | (Immelmann, 1966) |
| Passeriformes | Artamidae | Great woodswallow | Artamus maximus | (Billerman et al., 2022) |
| Passeriformes | Artamidae | Pied butcherbird | Cracticus nigrogularis | (Milligan, 1905) |
| Passeriformes | Artamidae | Australian magpie | Gymnorhina tibicen | (Kramer, 1930) |
| Passeriformes | Callaeidae | Huia | Heteralocha acutirostris | (Buller, 1888) |
| Passeriformes | Callaeidae | Saddleback | Philesturnus carunculatus | (Billerman et al., 2022) |
| Passeriformes | Campephagidae | Bar-bellied Cuckooshrike | Coracina striata | (Billerman et al., 2022) |
| Passeriformes | Cardinalidae | Lazuli bunting | Passerina amoena | (Miller, 1939) |
| Passeriformes | Cardinalidae | Painted bunting | Passerina ciris | (Billerman et al., 2022) |
| Passeriformes | Cinclosomatidae | Chestnut-breasted Quail-thrush | Cinclosoma castaneothorax | (Billerman et al., 2022) |
| Passeriformes | Cinclosomatidae | Chestnut Quail-thrush | Cinclosoma castanotum | (Billerman et al., 2022) |
| Passeriformes | Cinclosomatidae | Spotted Quail-thrush | Cinclosoma punctatum | (Billerman et al., 2022) |
| Passeriformes | Cisticolidae | Common tailorbird | Orthotomus sutorius | (Touche and Rickett, 1905) |
| Passeriformes | Corcoracidae | White-winged Chough | Corcorax melanorhamphos | (Hobbs, 1971) |
| Passeriformes | Corcoracidae | Apostlebird | Struthidea cinerea | (Baldwin, 1974) |
| Passeriformes | Corvidae | Transvolcanic jay | Aphelocoma ultramarina | (Swarth, 1900; Clark, 1973; Billerman et al., 2022) |
| Passeriformes | Corvidae | White-throated Magpie-Jay | Calocitta formosa | (Clark, 1973) |
| Passeriformes | Corvidae | White-necked Raven | Corvus albicollis | (Clark, 1973) |
| Passeriformes | Corvidae | Northwestern crow | Corvus caurinus | (Billerman et al., 2022) |
| Passeriformes | Corvidae | Carrion crow | Corvus corone | (Finn, 1922) |
| Passeriformes | Corvidae | Rook | Corvus frugilegus | (Clark, 1973) |
| Passeriformes | Corvidae | Blue jay | Cyanocitta cristata | (Clark, 1973) |
| Passeriformes | Corvidae | Steller's jay | Cyanocitta stelleri | (Risdon, 1960; Skutch, 1967) |
| Passeriformes | Corvidae | Purplish jay | Cyanocorax cyanomelas | (Wetmore, 1926) |
| Passeriformes | Corvidae | Green jay | Cyanocorax luxuosus | (Boswall, 1983) |
| Passeriformes | Corvidae | Brown jay | Cyanocorax morio | (Skutch, 1954) |
| Passeriformes | Corvidae | Eurasian nutcracker | Nucifraga caryocatactes | (Witherby et al., 1943) |
| Passeriformes | Corvidae | Canada jay | Perisoreus canadensis | (Billerman et al., 2022) |
| Passeriformes | Corvidae | Canada jay | Perisoreus canadensis | (Ouellet, 1970) |
| Passeriformes | Corvidae | Yellow-billed Magpie | Pica nuttalli | (Linsdale, 1937) |

| | | | | |
|---|---|---|---|---|
| Passeriformes | Corvidae | Red billed cough | Pyrrhocorax pyrrhocorax | (Hunt, 1815) |
| Passeriformes | Corvidae | Stresemann's bush-crow | Zavattariornis stresemanni | (Billerman et al., 2022) |
| Passeriformes | Dicruridae | Fork-tailed drongo | Dicrurus adsimilis | (Ali, 1961) |
| Passeriformes | Dicruridae | Shining drongo | Dicrurus atripennis | (Billerman et al., 2022) |
| Passeriformes | Dicruridae | White-bellied drongo | Dicrurus caerulescens | (Kramer, 1930) |
| Passeriformes | Dicruridae | Tablas drongo | Dicrurus menagei | (Billerman et al., 2022) |
| Passeriformes | Dicruridae | Velvet-mantled Drongo | Dicrurus modestus | (Billerman et al., 2022) |
| Passeriformes | Dicruridae | Western Square-tailed Drongo | Dicrurus occidentalis | (Billerman et al., 2022) |
| Passeriformes | Dicruridae | Greater Racket-tailed | Dicrurus paradiseus | (Simmons, 1963) |
| Passeriformes | Dicruridae | Sharpe's drongo | Dicrurus sharpei | (Billerman et al., 2022) |
| Passeriformes | Estrildidae | Red-headed Parrotfinch | Erythrura cyaneovirens | (Billerman et al., 2022) |
| Passeriformes | Estrildidae | Gouldian finch | Erythrura gouldiae | (Billerman et al., 2022) |
| Passeriformes | Estrildidae | Pink-billed Parrotfinch | Erythrura kleinschmidti | (Billerman et al., 2022) |
| Passeriformes | Estrildidae | Royal parrotfinch | Erythrura regia | (Billerman et al., 2022) |
| Passeriformes | Estrildidae | Common waxbill | Estrilda astrild | (Immelmann and Immelmann, 1967; Billerman et al., 2022) |
| Passeriformes | Estrildidae | Black-crowned Waxbill | Estrilda nonnula | (Goodwin, 1963; Billerman et al., 2022) |
| Passeriformes | Estrildidae | Black-tailed Waxbill | Estrilda perreini | (Billerman et al., 2022) |
| Passeriformes | Estrildidae | Crimson-rumped Waxbill | Estrilda rhodopyga | (Billerman et al., 2022) |
| Passeriformes | Estrildidae | Black-rumped Waxbill | Estrilda troglodytes | (Harrison, 1962) |
| Passeriformes | Estrildidae | Indian silverbill | Euodice malabarica | (Immelmann and Immelmann, 1967) |
| Passeriformes | Estrildidae | Lavender waxbill | Glaucestrilda caerulescens | (Harrison, 1962; Hinze, 2000) |
| Passeriformes | Estrildidae | Gray-headed Munia | Lonchura caniceps | (Restall, 1996) |
| Passeriformes | Estrildidae | Magpie munia | Lonchura fringilloides | (Restall, 1996) |
| Passeriformes | Estrildidae | Black munia | Lonchura stygia | (Restall, 1996) |
| Passeriformes | Estrildidae | White-collared oliveback | Nesocharis ansorgei | (Clement, 1999) |
| Passeriformes | Estrildidae | Java sparrow | Padda oryzivora | (Billerman et al., 2022) |
| Passeriformes | Estrildidae | Bronze mannikin | Spermestes cucullata | (Immelmann and Immelmann, 1967) |
| Passeriformes | Estrildidae | Western bluebill | Spermophaga haematina | (Harrison, 1966) |
| Passeriformes | Estrildidae | Red-eared Firetail | Stagonopleura oculata | (Immelmann, 1966; Billerman et al., 2022) |
| Passeriformes | Estrildidae | Violet-eared Waxbill | Uraeginthus granatinus | (Billerman et al., 2022) |
| Passeriformes | Falcunculidae | Crested shrike-tit | Falcunculus frontatus | (Noske, 1985) |
| Passeriformes | Fringillidae | European goldfinch | Carduelis carduelis | (Newton, 1967) |
| Passeriformes | Fringillidae | European greenfinch | Carduelis chloris | (Newton, 1967; Billerman et al., 2022) |
| Passeriformes | Fringillidae | Citril finch | Carduelis citrinella | (Billerman et al., 2022) |
| Passeriformes | Fringillidae | Common redpoll | Carduelis flammea | (Newton, 1967) |
| Passeriformes | Fringillidae | Hoary redpoll | Carduelis hornemanni | (Billerman et al., 2022) |
| Passeriformes | Fringillidae | Eurasian siskin | Carduelis spinus | (Newton, 1967) |
| Passeriformes | Fringillidae | American goldfinch | Carduelis tristis | (Coutlee, 1963; Billerman et al., 2022) |
| Passeriformes | Fringillidae | Common chaffinch | Fringilla coelebs | (Marler, 1956; Kear, 1962) |
| Passeriformes | Fringillidae | Oahu amakihi | Hemignathus flava | (Billerman et al., 2022) |
| Passeriformes | Fringillidae | Akiapolaau | Hemignathus munroi | (Billerman et al., 2022) |
| Passeriformes | Fringillidae | Kauai amakihi | Hemignathus stejnegeri | (Billerman et al., 2022) |
| Passeriformes | Fringillidae | Hawaii amakihi | Hemignathus virens | (Billerman et al., 2022) |
| Passeriformes | Fringillidae | Laysan ‘apapane | Himatione fraithii | (Fisher, 1903) |
| Passeriformes | Fringillidae | Apapane | Himatione sanguinea | (Billerman et al., 2022) |
| Passeriformes | Fringillidae | Twite | Linaria flavirostris | (Kear, 1962) |
| Passeriformes | Fringillidae | Red crossbill | Loxia curvirostra | (Newton, 1967; Billerman et al., 2022) |
| Passeriformes | Fringillidae | White-winged Crossbill | Loxia leucoptera | (Bent and Austin, 1968) |
| Passeriformes | Fringillidae | Parrot crossbill | Loxia pytyopsittacus | (Billerman et al., 2022) |
| Passeriformes | Fringillidae | Maui alauahio | Paroreomyza montana | (Billerman et al., 2022) |
| Passeriformes | Fringillidae | Maui parrotbill | Pseudonestor xanthophrys | (Billerman et al., 2022) |
| Passeriformes | Fringillidae | Island canary | Serinus canaria | (Billerman et al., 2022) |
| Passeriformes | Fringillidae | Streaky-headed Seedeater | Serinus gularis | (Billerman et al., 2022) |
| Passeriformes | Fringillidae | European serin | Serinus serinus | (Billerman et al., 2022) |
| Passeriformes | Fringillidae | Pine siskin | Spinus pinus | (Clark, 1973) |
| Passeriformes | Furnariidae | Pink-legged Graveteiro | Acrobatornis fonsecai | (Pacheco et al., 1996) |
| Passeriformes | Furnariidae | Olive-backed foliage-gleaner | Automolus infuscatus | (Zimmer, 2002) |
| Passeriformes | Furnariidae | Buff-throated Foliage-gleaner | Automolus ochrolaemus | (Skutch, 1969) |
| Passeriformes | Furnariidae | Rufous-fronted thornbird | Phacellodomus rufifrons | (Skutch, 1969) |
| Passeriformes | Furnariidae | Rufous-rumped foliage-gleaner | Philydor erythrocercum | (Rosenberg, 1997) |
| Passeriformes | Furnariidae | Russet-mantled Foliage-gleaner | Syndactyla dimidiata | (Robbins and Zimmer, 2005) |
| Passeriformes | Icteridae | Bay-winged Cowbird | Agelaioides badius | (Billerman et al., 2022) |
| Passeriformes | Icteridae | Yellow-shouldered Blackbird | Agelaius xanthomus | (Billerman et al., 2022) |

| | | | | |
|---|---|---|---|---|
| Passeriformes | Icteridae | Pale-eyed Blackbird | Agelaius xanthophthalmus | (Orians and Orians, 2000) |
| Passeriformes | Icteridae | Yellow-billed Cacique | Amblycercus holosericeus | (Skutch, 1967) |
| Passeriformes | Icteridae | Scarlet-rumped cacique | Cacicus uropygialis | (Skutch and Club, 1972) |
| Passeriformes | Icteridae | Melodious blackbird | Dives dives | (Skutch, 1954) |
| Passeriformes | Icteridae | Bobolink | Dolichonyx oryzivorus | (Gosse et al., 1847) |
| Passeriformes | Icteridae | Rusty blackbird | Euphagus carolinus | (Billerman et al., 2022) |
| Passeriformes | Icteridae | Brewer's blackbird | Euphagus cyanocephalus | (La Rivers, 1941; Billerman et al., 2022) |
| Passeriformes | Icteridae | Baltimore oriole | Icterus galbula | (Wellman, 1928; Billerman et al., 2022) |
| Passeriformes | Icteridae | Baltimore oriole | Icterus galbula | (Wellman, 1928) |
| Passeriformes | Icteridae | Montserrat oriole | Icterus oberi | (Billerman et al., 2022) |
| Passeriformes | Icteridae | Orchard oriole | Icterus spurius | (Lohrer, 1977; Billerman et al., 2022) |
| Passeriformes | Icteridae | Brown-headed Cowbird | Molothrus ater | (Bent and Austin, 1968) |
| Passeriformes | Icteridae | Boat-tailed Grackle | Quiscalus major | (Billerman et al., 2022) |
| Passeriformes | Icteridae | Great-tailed Grackle | Quiscalus mexicanus | (Clark, 1973) |
| Passeriformes | Icteridae | Common grackle | Quiscalus quiscula | (Roberts, 1932; Clark, 1973) |
| Passeriformes | Icteriidae | Yellow-breasted Chat | Icteria virens | (Ficken, 1962; Billerman et al., 2022) |
| Passeriformes | Laniidae | Commonfiscal | Lanius collaris | (Cooper, 1971) |
| Passeriformes | Laniidae | Fiscal shrike | Lanius collaris | (Van Someren, 1956) |
| Passeriformes | Laniidae | Red-backed Shrike - | Lanius collurio | (Ash, 1970) |
| Passeriformes | Laniidae | Great grey shrike | Lanius excubitor | (Cade, 1967; Billerman et al., 2022) |
| Passeriformes | Laniidae | Loggerhead shrikes | Lanius ludovicianus | (Miller, 1931) |
| Passeriformes | Laniidae | Lesser gray shrike | Lanius minor | (Ullrich, 1971) |
| Passeriformes | Leiothrichidae | Blue-winged minla | Actinodura cyanouroptera | (Finn, 1908) |
| Passeriformes | Leiothrichidae | Striated babbler | Argya earlei | (Finn, 1908) |
| Passeriformes | Leiothrichidae | Black-headed sibia | Heterophasia desgodinsi | (Finn, 1908) |
| Passeriformes | Leiothrichidae | Silver-eared mesia | Leiothrix argentauris | (Finn, 1908; Gibson, 1991) |
| Passeriformes | Leiothrichidae | Red-billed Leiothrix | Leiothrix lutea | (Kramer, 1930) |
| Passeriformes | Leiothrichidae | White-throated Laughingthrush | Pterorhinus albogularis | (Ali, 1961) |
| Passeriformes | Leiotrichidae | Chinese hwamei | Garrulax canorus | (Kramer, 1930) |
| Passeriformes | Leiotrichidae | White-crested Laughingthrush | Garrulax leucolophus | (Kramer, 1930) |
| Passeriformes | Leiotrichidae | Striated laughingthrush | Grammatoptila striata | (Kramer, 1930) |
| Passeriformes | Leiotrichidae | Yellow-billed Babbler | Turdoides affinis | (Ali, 1961) |
| Passeriformes | Leiotrichidae | Arrow-marked Babbler | Turdoides jardineii | (Billerman et al., 2022) |
| Passeriformes | Locustellidae | New zealand fernbird | Poodytes punctatus | (Best, 1979) |
| Passeriformes | Malaconotidae | Ethiopian boubou | Laniarius aethiopicus | (Billerman et al., 2022) |
| Passeriformes | Malaconotidae | Grey-headed bushshrike | Malaconotus blanchoti | (Billerman et al., 2022) |
| Passeriformes | Malaconotidae | Fiery-breasted Bushshrike | Malaconotus cruentus | (Billerman et al., 2022) |
| Passeriformes | Malaconotidae | Brubru | Nilaus afer | (Billerman et al., 2022) |
| Passeriformes | Malacotonidae | The brown-crowned tchagra | Tchagra australis | (Van Someren, 1956) |
| Passeriformes | Meliphagidae | Long-billed honeyeater | Melilestes megarhynchus | (Brown and Hopkins, 2002) |
| Passeriformes | Meliphagidae | Mimic meliohaga | Meliphaga analoga | (Brown and Hopkins, 2002) |
| Passeriformes | Meliphagidae | Puff-backed honeyeater | Meliphaga aruensis | (Brown and Hopkins, 2002) |
| Passeriformes | Meliphagidae | Mottle-breasted honeyeater | Meliphaga mimikae | (Brown and Hopkins, 2002) |
| Passeriformes | Meliphagidae | Helmeted friarbird | Philemon buceroides | (Brown and Hopkins, 2002) |
| Passeriformes | Meliphagidae | Striped honeyeater | Plectorhyncha lanceolata | (Billerman et al., 2022) |
| Passeriformes | Meliphagidae | Tawny-breasted Honeyeater | Xanthotis flaviventer | (Brown and Hopkins, 2002) |
| Passeriformes | Meliphagidae | Spotted honeyeater | Xanthotis polygrammus | (Brown and Hopkins, 2002) |
| Passeriformes | Menuridae | Superb lyrebird | Menura novaehollandie | (Austin et al., 2019) |
| Passeriformes | Mimidae | California thrasher | Toxostoma redivivum | (Clark, 1973) |
| Passeriformes | Mohouidae | Yellowhead | Mohoua ochrocephala | (Billerman et al., 2022) |
| Passeriformes | Monarchidae | Elepaio | Chasiempis sandwichensis | (Billerman et al., 2022) |
| Passeriformes | Monarchidae | Black-naped Monarch | Hypothymis azurea | (Billerman et al., 2022) |
| Passeriformes | Monarchidae | Black-faced Monarch | Monarcha melanopsis | (Harrison, 1969) |
| Passeriformes | Motacillidae | White wagtail | Motacilla alba | (Billerman et al., 2022) |
| Passeriformes | Nectariniidae | Seychelles sunbird | Cinnyris dussumieri | (Greig-Smith, 1980) |
| Passeriformes | Neosittidae | Varied sittella | Daphoenositta chrysoptera | (Billerman et al., 2022) |
| Passeriformes | Panuridae | Bearded reedling | Panurus biarmicus | (Koenig, 1952) |
| Passeriformes | Paradisaeidae | Magnificent Bird of Paradise | Cicinnurus magnificus | (Beehler and Dumbacher, 1996; Brown and Hopkins, 2002) |
| Passeriformes | Paradisaeidae | Crinkle-collared manucode | Manucodia chalybatus | (Brown and Hopkins, 2002) |

| | | | | |
|---|---|---|---|---|
| Passeriformes | Paradisaeidae | Raggiana bird-of-paradise | Paradisaea raggiana | (Beehler and Dumbacher, 1996; Brown and Hopkins, 2002) |
| Passeriformes | Paradisaeidae | Red bird-of-paradise | Paradisaea rubra | (Frith, 1976b) |
| Passeriformes | Paradisaeidae | Wahnes's parotia | Parotia wahnesi | (Frith and Frith, 1979) |
| Passeriformes | Paradisaeidae | Magnificent riflebird | Ptiloris magnificus | (Beehler and Dumbacher, 1996; Brown and Hopkins, 2002) |
| Passeriformes | Paradisaeidae | Twelve-wired bird-of-paradise | Seleucidis melanoleucus | (Rand and Gilliard, 1967) |
| Passeriformes | Paradoxornithidae | Yellow-eyed Babbler | Chrysomma sinense | (Kramer, 1930) |
| Passeriformes | Paradoxornithidae | The yellow-eyed babbler | Chrysomma sinense | (Harper, 1901) |
| Passeriformes | Paradoxornithidae | Gray-headed Parrotbill | Paradoxornis gularis | (Billerman et al., 2022) |
| Passeriformes | Paridae | Oak titmouse | Baeolophus inornatus | (Root, 1967) |
| Passeriformes | Paridae | Fire-capped Tit | Cephalopyrus flammiceps | (Billerman et al., 2022) |
| Passeriformes | Paridae | Gray tit | Parus afer | (Yince, 1964) |
| Passeriformes | Paridae | Black-capped Chickadee | Parus atricapillus | (Brewer, 1961; Clark, 1973) |
| Passeriformes | Paridae | Eurasian blue tit | Parus caeruleus | (Yince, 1964) |
| Passeriformes | Paridae | Carolina chickadee | Parus carolinensis | (Brewer, 1961) |
| Passeriformes | Paridae | Mountain chickadee | Parus gambeli | (Clark, 1973) |
| Passeriformes | Paridae | Sombre tit | Parus lugubris | (Löhrl, 1966) |
| Passeriformes | Paridae | Japanese tit | Parus major | (Yince, 1964) |
| Passeriformes | Paridae | Carolina chickadee | Poecile carolinensis | (Brewer, 1961) |
| Passeriformes | Parulidae | Yellow-rumped Warbler | Dendroica coronata | (Conway et al., 2002) |
| Passeriformes | Parulidae | Worm-eating Warbler | Helmitheros vermivorum | (Billerman et al., 2022) |
| Passeriformes | Passerellidae | Grassland sparrow | Ammodramus humeralis | (Billerman et al., 2022) |
| Passeriformes | Passeridae | House sparrow | Passer domesticus | (Summers-Smith, 1963) |
| Passeriformes | Platysteiridae | Chinspot batis | Batis molitor | (Harris, 2010) |
| Passeriformes | Ploceidae | Aldabra fody | Foudia aldabrana | (Frith, 1976a) |
| Passeriformes | Ploceidae | Comoros fody | Foudia eminentissima | (Cheke and Diamond, 1986) |
| Passeriformes | Ploceidae | Red-headed Malimbe | Malimbus rubricollis | (Billerman et al., 2022) |
| Passeriformes | Ploceidae | Baya weaver | Ploceus philippinus | (Das et al., 2015) |
| Passeriformes | Pomatostomidae | Papuan babbler | Pomatostomus isidorei | (Billerman et al., 2022) |
| Passeriformes | Psophodidae | Eastern whipbird | Psophodes olivaceus | (Billerman et al., 2022) |
| Passeriformes | Pycnonotidae | Yellow-eared Bulbul | Pycnonotus penicillatus | (Chandrasiri and Mahaulpatha, 2019) |
| Passeriformes | Regulidae | Ruby-crowned kinglet | Regulus calendula | (Ross, 1924) |
| Passeriformes | Regulidae | The tenerife kinglet | Regulus teneriffae | (Löhrl et al., 1996) |
| Passeriformes | Remizidae | Southern penduline-tit | Anthoscopus minutus | (Skead, 1959; Billerman et al., 2022) |
| Passeriformes | Remizidae | Verdin | Auriparus flaviceps | (Taylor, 1971) |
| Passeriformes | Remizidae | Eurasian penduline-tit | Remiz pendulinus | (Billerman et al., 2022) |
| Passeriformes | Rhipiduridae | New zealand fantail | Rhipidura fuliginosa | (Harrison, 1969) |
| Passeriformes | Rhipiduridae | Willie-wagtail | Rhipidura leucophrys | (Koenig, 1952; Billerman et al., 2022) |
| Passeriformes | Rhipiduridae | Rufous fantail | Rhipidura rufifrons | (Billerman et al., 2022) |
| Passeriformes | Sittidae | Snowy-browed Nuthatch | Sitta villosa | (Billerman et al., 2022) |
| Passeriformes | Sturnidae | Brahminy starling | Sturnus pagodarum | (Bhardwaj and Kumar, 2004) |
| Passeriformes | Sylviidae | Wrentit | Chamaea fasciata | (Billerman et al., 2022) |
| Passeriformes | Sylviidae | Wrentit | Chamaea fasciata | (Erickson, 1938) |
| Passeriformes | Thraupidae | Woodpecker finch | Camarhynchus pallidus | (Millikan, 1967) |
| Passeriformes | Thraupidae | Small tree-finch | Camarhynchus parvulus | (Bowman, 1961) |
| Passeriformes | Thraupidae | Large Tree-finch | Camarhynchus psittacula | (Bowman, 1961) |
| Passeriformes | Thraupidae | Green warbler finch | Certhidea olivacea | (Bowman, 1961) |
| Passeriformes | Thraupidae | Lesser antillean bullfinch | Loxigilla noctis | (Billerman et al., 2022) |
| Passeriformes | Thraupidae | Puerto rican bullfinch | Loxigilla portoricensis | (Billerman et al., 2022) |
| Passeriformes | Thraupidae | Fawn-breasted tanager | Pipraeidea melanonota | (Brown and Neto, 1976) |
| Passeriformes | Thraupidae | Temminck's seedeater | Sporophila falcirostris | (Areta et al., 2013) |
| Passeriformes | Thraupidae | Buffy-fronted Seedeater | Sporophila frontalis | (Areta et al., 2013) |
| Passeriformes | Thraupidae | Yellow-faced Grassquit | Tiaris olivaceus | (Baptista, 1976) |
| Passeriformes | Timaliidae | White-browed Scimitar-Babbler | Pomatorhinus schisticeps | (Kramer, 1930) |
| Passeriformes | Turdidae | Wood thrush | Hylocichla mustelina | (Reed, 1905) |
| Passeriformes | Turdidae | Townsend's solitaire | Myadestes townsendi | (Billerman et al., 2022) |
| Passeriformes | Tyrannidae | Pacific-slope Flycatcher | Empidonax difficilis | (Bent, 1942) |
| Passeriformes | Tyrannidae | Willow flycatcher | Empidonax traillii | (Brewster, 1865; La Rivers, 1941) |
| Passeriformes | Tyrannidae | Streaked flycatcher | Myiodynastes maculatus | (Gross, 1950) |
| Passeriformes | Tyrannidae | White-rumped Monjita | Xolmis velatus | (Hudson, 1920) |
| Passeriformes | Vangidae | Red-shouldered Vanga | Calicalicus rufocarpalis | (Billerman et al., 2022) |
| Passeriformes | Vangidae | Helmet vanga | Euryceros prevostii | (Safford and Hawkins, 2020) |
| Passeriformes | Vangidae | Sickle-billed Vanga | Falculea palliata | (Billerman et al., 2022) |
| Passeriformes | Vangidae | Tylas vanga | Tylas eduardi | (Billerman et al., 2022) |
| Passeriformes | Viduidae | Pin-tailed Whydah | Vidua macroura | (Billerman et al., 2022) |
| Passeriformes | Vireonidae | Rufous-browed Peppershrike | Cyclarhis gujanensis | (Skutch, 1967) |
| Passeriformes | Vireonidae | Gray-eyed Greenlet | Hylophilus amaurocephalus | (Billerman et al., 2022) |

| | | | | |
|---|---|---|---|---|
| Passeriformes | Vireonidae | Bells vireo | Vireo bellii | (Nolan, 1960) |
| Passeriformes | Vireonidae | Noronha vireo | Vireo gracilirostris | (Billerman et al., 2022) |
| Passeriformes | Vireonidae | White-eyed Vireo | Vireo griseus | (Billerman et al., 2022) |
| Passeriformes | Vireonidae | Plumbeous vireo | Vireo plumbeus | (Billerman et al., 2022) |
| Passeriformes | Vireonidae | Blue-headed Vireo | Vireo solitarius | (Bent, 1965; Billerman et al., 2022) |
| Passeriformes | Vireonidae | Gray vireo | Vireo vicinior | (Billerman et al., 2022) |
| Passeriformes | Vireonidae | Chestnut-sided Shrike-Vireo | Vireolanius melitophrys | (Billerman et al., 2022) |
| Passeriformes | Zosteropidae | Mascarene Grey White-eye | Zosterops borbonica | (Gill, 1971) |
| Passeriformes | Zosteropidae | Réunion olive white-eye | Zosterops olivacea | (Gill, 1971) |
| Passeriformes | Zosteropidae | Swinhoe's white-eye | Zosterops simplex | (Finn, 1908) |
| Pelecaniformes | Ardeidae | Grey heron | Ardea cinerea | (Pistorius, 2008) |
| Pelecaniformes | Ardeidae | Blue heron | Ardea herodias | (Moseley, 1936) |
| Pelecaniformes | Ardeidae | Green-backed Heron | Ardeola striata | (Higuchi, 1986) |
| Pelecaniformes | Ardeidae | Pacific reef heron | Egretta sacra | (Beckmann, 2008) |
| Pelecaniformes | Threskiornithidae | Australian white ibis | Threskiornis molucca | (Morris, 1973) |
| Piciformes | Capitonidae | Red-headed Barbet | Eubucco bourcierii | (Billerman et al., 2022) |
| Piciformes | Picidae | Golden-fronted woodpecker | Melanerpes aurifrons | (Martin and Kroll, 1975) |
| Piciformes | Ramphastidae | Plate-billed Mountain-Toucan | Andigena laminirostris | (Billerman et al., 2022) |
| Piciformes | Ramphastidae | Chestnut-mandibled toucan | Ramphastos ambiguus | (Skutch and Club, 1972) |
| Piciformes | Semnornithidae | Prong-billed Barbet | Semnornis frantzii | (Billerman et al., 2022) |
| Piciformes | Semnornithidae | Toucan barbet | Semnornis ramphastinus | (Billerman et al., 2022) |
| Psittaciformes | Cacatuidae | Tanimbar corella | Cacatua goffiniana | (Auersperg et al., 2014) |
| Psittaciformes | Cacatuidae | Salmon-crested cockatoo | Cacatua moluccensis | (Boswall, 1983) |
| Psittaciformes | Cacatuidae | Palm cockatoo | Probosciger aterrimus | (Wallace, 1869) |
| Psittaciformes | Psittacidae | Hyacinth macaws | Anodorhynchus hyacinthinus | (Borsari and Ottoni, 2005) |
| Psittaciformes | Psittacidae | Cape parrot | Poicephalus robustus | (Dowsett-Lemaire, 2004) |
| Psittaciformes | Strigopidae | Kakapo | Strigops habroptilus | (Best, 1984) |
| Psittasiformes | Psittaculidae | Australian ringneck | Barnardius zonarius | (Milligan, 1905; Nichols, 1978) |
| Strigiformes | Strigidae | Long-eared owl | Asio otus | (MacGillivray) |
| Strigiformes | Strigidae | Akun Eagle-owl | Bubo leucostictus | (Billerman et al., 2022) |
| Strigiformes | Strigidae | Whiskered Screech-owl | Megascops trichopsis | (Billerman et al., 2022) |
| Strigiformes | Tytonidae | Barn owl | Tyto alba | (Csermely and Gaibani, 1998) |
| Suliformes | Phalacrocoracidae | Reed cormorant | Microcarbo africanus | (Olver, 1984) |
| Suliformes | Phalacrocoracidae | Little pied cormorant | Microcarbo melanoleucos | (Cole, 1908) |
| Suliformes | Phalacrocoracidae | Little cormorant | Microcarbo niger | (Cole, 1908) |

MacGillivray W. A history of British birds, indigenous and migratory: including their organization, habits, and relations; remarks on classification and nomenclature; an account of the principal organs of birds, and observations relative to practical ornithology ... London, Printed for Scott, Webster, and Geary, 1837-1852. Available from: https://www.biodiversitylibrary.org/item/33142

Marler P. 1956. Behaviour of the Chaffinch Fringilla Coelebs. Behaviour Supplement:III–184.

Martin JW, Kroll JC. 1975. Hoarding of corn by Golden-fronted Woodpeckers. The Wilson Bulletin 87:553–553.

Miller AH. 1931. Systematic revision and natural history of the American shrikes (Lanius). University of California Press.

Miller AH. 1939. Foraging Dexterity of a Lazuli Bunting. The Condor 41:255–256.

Milligan AlexWm. 1905. Notes on a Trip to the Yandanooka District, Western Australia. Emu - Austral Ornithology 4:151–157.

Millikan GC. 1967. Observations on Galápagos tool-using finches in captivity. Living Bird 6:23–41.

Morris AK. 1973. White Ibis eats snake. EMU 73:73–73.

Moseley EL. 1936. Blue Heron Colonies in Northern Ohio. The Wilson Bulletin 48:3–11.

Mundy PJ, Robertson AS, Komen J, O’connor TG. 1986. Attacks by black eagles on vultures. Raptor research 20:61–64.

Newton I. 1967. The Adaptive Radiation and Feeding Ecology of Some British Finches. Ibis 109:33–96.

Nichols OG. 1978. From Field and Study: The Port Lincoln Parrot (Barnardius zonarius zonarius) feeding on Lerps at Kalgoorlie. The Western Australian Naturalist 14:76–77.

Nolan V. 1960. Breeding Behavior of the Bell Vireo in Southern Indiana. The Condor 62:225–244.

Noske RA. 1985. Left-footedness and tool-using in the varied sittella Daphoenositta chrysoptera and crested shrike-tit Falcunculus frantatus. Corella 9:63–64.

Olver MD. 1984. Breeding Biology of the Reed Cormorant. Ostrich 55:133–140.

Orians GH, Orians EN. 2000. Observations of the Pale-Eyed Blackbird in Southeastern Peru. The Condor 102:956–958.

Oswald T, Curtice B, Bolander M, Lopez C. 2023. Observation of Claw Use and Feeding Behavior of the Red-Legged Seriema and Its Implication for Claw Use in Deinonychosaurs. jana 50:17–21.

Ouellet H. 1970. Further observations on the food and predatory habits of the gray jay. Can J Zool 48:327–330.

Pacheco JF, Whitney BM, Gonzaga LP. 1996. A new genus and species of furnariid (Aves: furnariidae) from the cocoa-growing region of southeastern Bahia, Brazil. Wilson Bulletin 108:397–434.

Pistorius PA. 2008. Grey Heron (Ardea cinerea) Predation on the Aldabra White-throated Rail (Dryolimnas cuvieri aldabranus). wils 120:631–632.

Pruett-Jones SG. 1980. Team-Hunting and Food Sharing in Parasitic Jaegers. The Wilson Bulletin 92:524–526.

Rand AL, Gilliard ET. 1967. Handbook of new guinea birds. Weidenfeld & Nicolson.

Reed Chester A (Chester Albert). 1905. American ornithology, for home and school. [Worcester, Mass, Charles K. Reed], 1901-[1906]. Available from: https://www.biodiversitylibrary.org/item/101521

Restall R. 1996. Munias and mannikins. Sussex: Pica Press.

Risdon DHS. 1960. The breeding of Steller’s Jay ( Cyanocitta stelleri) at Dudley Zoo. In: The Avicultural magazine. Vol. v.66 (1960). Ascot, Berkshire, etc, Avicultural Society, etc,. Available from: https://www.biodiversitylibrary.org/item/253794

Robbins MB, Zimmer KJ. 2005. Taxonomy, vocalisations and natural history of Philydor dimidiatum (Furnariidae), with comments on the systematics of Syndactyla and Simoxenops. Bulletin of the British Ornithologists’ Club 125:212–228.

Roberts TS. 1932. The birds of Minnesota. Vol. II. Minneapolis: Univ. of Minnesota Press.

Root RB. 1967. The Niche Exploitation Pattern of the Blue-Gray Gnatcatcher. Ecological Monographs 37:317–350.

Rosenberg KV. 1997. Ecology of Dead-Leaf Foraging Specialists and Their Contribution to Amazonian Bird Diversity. Ornithological Monographs:673–700.

Ross RC. 1924. Occurrence and Behavior of Certain Shorebirds in Southern California. The Condor 26:90–92.

Rowan M. 1967. A study of the colies of southern Africa. Ostrich 38:63–115.

Safford R, Hawkins F. 2020. The birds of Africa: Volume VIII: The Malagasy region: Madagascar, Seychelles, Comoros, Mascarenes. Bloomsbury Publishing.

Seavy NE, Schulze MD, Whitacre DF (David F, Vasquez MA. 1998. Breeding Biology and Behavior of the Plumbeous Kite. The Wilson bulletin 110:77–85.

Simmons K. 1963. Some behaviour characters of the babblers (Timaliidae). Avicultural Magazine 69:183–193.

Skead C. 1959. A study of the Cape Penduline Tit Anthoscopus minutus minutus (Shaw & Nodder). Ostrich 30:274–288.

Skutch A. 1954. Life histories of Central American birds, vol 1 (Pacific Coast Avifauna 31). Cooper Ornithological Society, Berkeley.

Skutch AF. 1967. Life histories of Central American highland birds. The Club,. Available from: https://www.biodiversitylibrary.org/item/260537

Skutch AF. 1969. A Study of the Rufous-Fronted Thornbird and Associated Birds: Part 1. Life History of the Rufous-Fronted Thornbird. The Wilson Bulletin 81:5–43.

Skutch Alexander F (Alexander Frank), Club NO. 1972. Studies of tropical American birds. Cambridge, Mass, The Club, 1972. Available from: https://www.biodiversitylibrary.org/item/259940

Smith G. 1971. The use of the foot in feeding, with especial reference to parrots. Avicult Mag 77:93–100.

Summers-Smith D. 1963. The House Sparrow. London.

Sustaita D, Gloumakov Y, Tsang LR, Dollar AM. 2019. Behavioral correlates of semi-zygodactyly in Ospreys (Pandion haliaetus) based on analysis of internet images. PeerJ 7:e6243.

Swarth HS. 1900. Birds of the Huachuca Mountains, Arizona. The Club.

Taylor WK. 1971. A Breeding Biology Study of the Verdin, Auriparus flaviceps (Sundevall) in Arizona. The American Midland Naturalist 85:289–328.

Thomsett S. 1981. Some observations on the bat hawk Macheiramphus alcinus. Scopus 5:56–56.

Touche JDL, Rickett CB. 1905. II.-Further Notes on the Nesting o j Birds in the Province of Foibkien, S.E. China. Ibis 47:25–67.

Ullrich B. 1971. Untersuchungen zur Ethologie und Ökologie des Rotkopfwürgers (Lanius senator) in Südwestdeutschland im Vergleich zu Raubwürger (L. excubitor), Schwarzstirnwürger (L. minor) und Neuntöter (L. collurio).

Van Someren VGL. 1956. Days with Birds: Studies of Habits of some East Africa birds. In: Fieldiana Zoology. Vol. v.38 (1956). [Chicago] Chicago Natural History Museum [1945?-. Available from: https://www.biodiversitylibrary.org/item/21311

Wallace, AR. 1869. The Malay Archipelago: The land of the orang-utan, and the bird of paradise. A narrative of travel, with studies of man and nature. London: Macmillan and Co. Available from: http://darwin-online.org.uk/converted/Ancillary/1869_MalayArchipelago_A1013.1/1869_MalayArchipelago_A1013.1.1.html

Wellman GB. 1928. Baltimore Oriole Feeding on Larvae of Needle Miner. The Auk 45:507–507.

Wetmore A. 1926. Observations on the birds of Argentina, Paraguay, Uruguay, and Chile. US Government Printing Office.

Witherby HF, 1873-1944, Jourdain FCR, 1865-1940, Ticehurst NF, b. 1873, Tucker BW, 1901-. 1943. Handbook of British birds. H.F. & G. Witherby ltd. Available from: https://scholar.google.com/scholar_lookup?title=handbook+of+British+birds&author=Witherby%2C+H.+F.+%28Harry+Forbes%29&publication_year=1943

Yince MA. 1964. Use of the Feet in Feeding by the Great Tit Parus Major. Ibis 106:508–529.

Zimmer KJ. 2002. Species Limits in Olive-Backed Foliage-Gleaners (Automolus: Furnariidae). The Wilson bulletin 114:20–37.
